## Supplementary material for "An individualized causal framework for learning intercellular communication networks that define microenvironments of individual tumors": A compressed folder containing supplementary tables: Table S7 Network_cl_clinical_table.docx

**Table S7. Distribution of clinical features among different network-based cluster subtypes**

|  | **network_cl1** | **network_cl2** | **network_cl3** | **network_cl4** |
| --- | --- | --- | --- | --- |
| **Age** | 59.18182±12.04 | 58.49±13.52 | 62.67±11.26 | 61.37±10.77 |
| **Staging** |  |  |  |  |
| I | 5 | 2 | 10 | 3 |
| II | 18 | 19 | 39 | 22 |
| III | 15 | 25 | 40 | 25 |
| IVA | 48 | 55 | 91 | 70 |
| IVB | 1 | 1 | 6 | 3 |
| IVC | 0 | 4 | 3 | 0 |
| **Node Invasion**  No | 13 | 8 | 21 | 12 |
| Yes | 58 | 98 | 161 | 105 |
| **Path M** |  |  |  |  |
| M0 | 86 | 100 | 181 | 121 |
| M1 | 0 | 3 | 3 | 0 |
| MX | 2 | 5 | 9 | 4 |
| **Path N** |  |  |  |  |
| N0 | 36 | 46 | 98 | 63 |
| N1 | 11 | 22 | 30 | 20 |
| N2 | 3 | 8 | 5 | 3 |
| N2a | 4 | 1 | 7 | 5 |
| N2b | 22 | 15 | 25 | 19 |
| N2c | 8 | 9 | 19 | 9 |
| N3 | 1 | 1 | 4 | 3 |
| NX | 3 | 6 | 5 | 4 |
| Path T |  |  |  |  |
| T1 | 10 | 6 | 14 | 5 |
| T2 | 33 | 26 | 61 | 31 |
| T3 | 21 | 32 | 45 | 37 |
| T4 | 2 | 7 | 6 | 9 |
| T4a | 19 | 35 | 60 | 41 |
| T4b | 0 | 0 | 3 | 0 |
| TX | 3 | 2 | 4 | 3 |
| HPV_status^***^ |  |  |  |  |
| HPV- | 50 | 97 | 157 | 110 |
| HPV+ | 32 | 6 | 26 | 8 |
| Site |  |  |  |  |
| Alveolar Ridge^*^ | 1 | 7 | 8 | 2 |
| Base of tongue | 8 | 3 | 12 | 4 |
| Buccal Mucosa | 2 | 7 | 7 | 5 |
| Floor of mouth^**^ | 4 | 18 | 23 | 17 |
| Hypopharynx | 1 | 0 | 6 | 3 |
| Larynx^***^ | 16 | 19 | 36 | 45 |
| Lip | 1 | 0 | 1 | 1 |
| Oral Cavity^*^ | 12 | 14 | 28 | 19 |
| Oral Tongue^***^ | 19 | 37 | 54 | 20 |
| Tonsil^***^ | 24 | 3 | 10 | 6 |
| Hard Palate | 0 | 1 | 6 | 0 |
| Oropharynx | 0 | 1 | 4 | 4 |

* p-value < 0.05, ** p-values < 0.01, *** p < 0.001
